# 7,000 years of turnover: historical contingency and human niche construction shape the Caribbean’s Anthropocene biota

**DOI:** 10.1101/2020.03.05.978924

**Authors:** Melissa E. Kemp, Alexis M. Mychajliw, Jenna Wadman, Amy Goldberg

## Abstract

The human-mediated movement of species across biogeographic boundaries—whether intentional or accidental—is dramatically reshaping the modern world. Yet, humans have been reshaping ecosystems and translocating species for millennia, and acknowledging the deeper roots of these phenomena is important for contextualizing present-day biodiversity loss, ecosystem functioning, and management needs. Here, we present the first database of terrestrial vertebrate species introductions spanning the entire anthropogenic history of a system: the Caribbean. We employ this ~7,000 year dataset to assess the roles of historical contingency and priority effects in shaping present-day community structure and conservation outcomes, finding that serial human colonization events contributed to habitat modifications and species extinctions that shaped the trajectories of subsequent species introductions by other human groups. We contextualized spatial and temporal patterns of species introductions within cultural practices and population histories of Indigenous, colonial, and modern human societies, and show that the taxonomic and biogeographic diversity of introduced species reflects diversifying reasons for species introductions through time. Recognition of the complex social and economic structures across the 7,000-year human history of the Caribbean provides the necessary context for interpreting the formation of an Anthropocene biota.

## Introduction

A wide range of disciplines continue to debate the origin and defining characteristics of the Anthropocene [1]. Yet, an undeniable signal of global human impact is the formation of novel ecosystems and non-analogue assemblages, which have been created by the intentional and accidental movement of species outside their native ranges in the context of land use change and selective extinctions [2]. Human-mediated species translocations have fundamentally changed the planet’s biogeographic assembly rules by eroding the barriers separating islands, continents, and hemispheres [3]. This reshaping of species ranges is not unique to present day human societies: today’s introduced species are part of a much longer legacy, with the earliest documented translocation occurring approximately 20,000 years ago [4]. Since this Late Pleistocene beginning, the advent of agricultural domestication and advancement of maritime technology vastly increased the number of translocations and the geographic distance of the translocation in the Holocene [5], and subsequently, colonial trade, industrial shipping, and the wide-ranging effects of globalization have generated the Anthropocene biota that define our modern landscapes [6].

The ecological and economic impacts of such translocated species today are intensively studied in management contexts, but whether and how the long-term (millennial-scale) sequence of human-mediated species introductions have hindered or promoted subsequent introductions and the establishment of these species remains poorly resolved. As priority effects – the order in which species arrive in a community – seem to favor the success of non-native species [7], the incorporation of historical contingency and an understanding of deeper time translocations can guide new research directions that help invasive species management while also provoking new eco-evolutionary questions. This approach also recognizes the agency of a diverse set of indigenous groups in shaping landscapes prior to subsequent colonizations, as restoration baselines in the western hemisphere are generally biased to the arrival of Columbus and do not incorporate pre-European baseline shifts [8].

The insular Caribbean is an exemplary replicate system in which to study the accumulation of non-native species over time and their ecological consequences, as it is a biodiversity hotspot that has undergone multiple distinct waves of human colonization and was the center of the Columbian Exchange. This region, situated between North and South America, encompasses over 7,000 islands comprising a range of sizes and geologic formations that fall into three major biogeographic groupings: the Greater Antilles, the Bahamas, and the Lesser Antilles (Figure 1). The combination of islands within and between these groupings has yielded an exceptionally biodiverse biota [9], though this diversity has been recently pruned: today’s 73 mammal species (13 nonvolant mammals and 60 bat species; [10]) represent a small remnant of the 130 species once found during the Holocene. Because many different cultural groups from around the globe have colonized the Caribbean over time, it is possible to compare trace how biotas that may start out similar become increasingly disparate due to extinction and introduction dynamics related to economics, trade, and culture.

**Figure 1.**
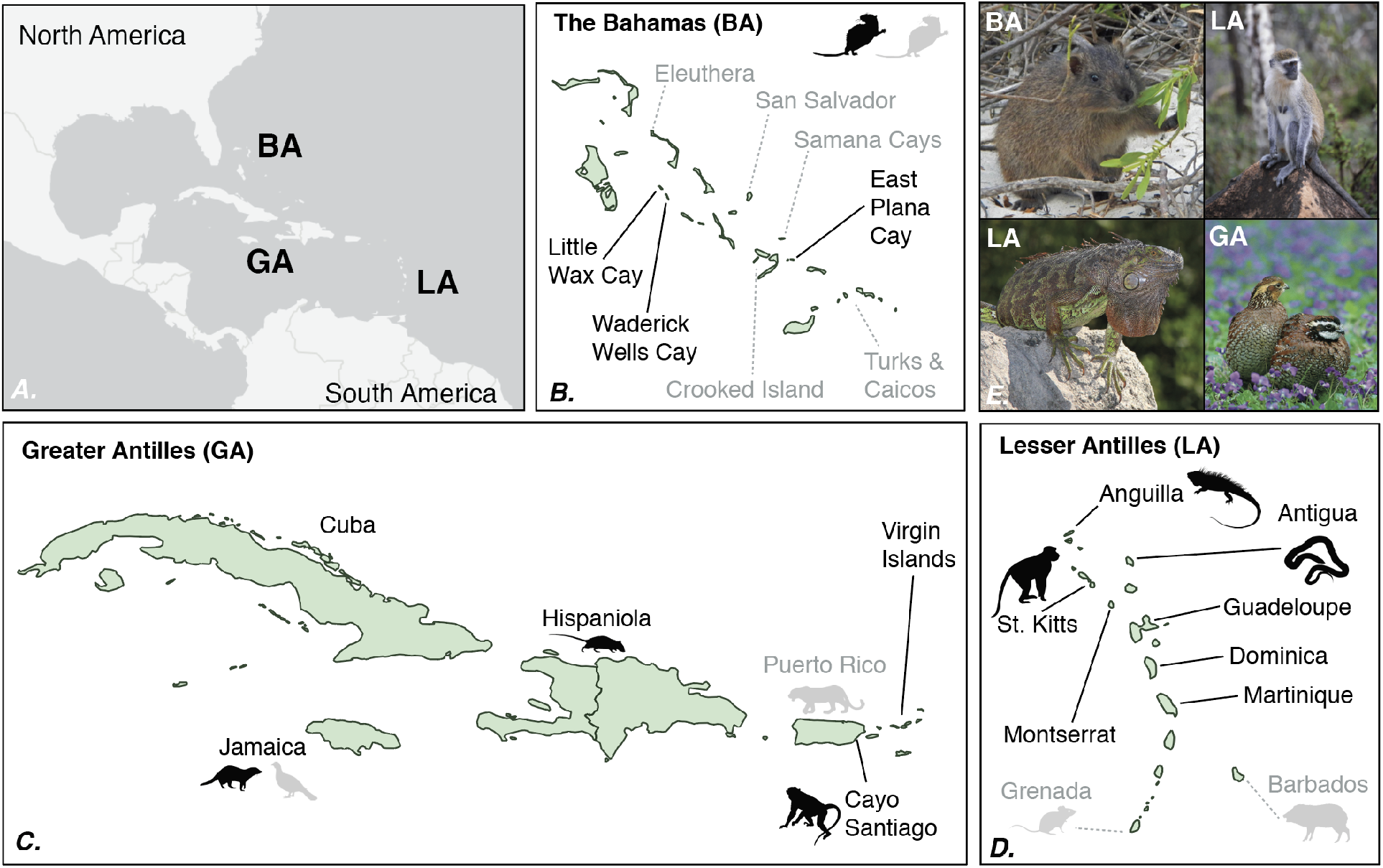
*The Caribbean and select species introductions. A. Relative locations of the Bahamas (BA), Greater Antilles (GA), and Lesser Antilles (LA). For B.-D., black symbols represent extant populations, and gray symbols represent failed introductions*. *B. Bahamian hutias (*Geocapromys ingrahami, E. BA) *were translocated by Late Ceramic people across the archipelago (gray, dashed lines) [11] and in 1973, extant populations (black lines) were moved for conservation*. *C. The bobwhite (*Colinus virgianus, E. GA) *was introduced to colonial Jamaica in 1747 for game, then extirpated with the 1872 introduction of the mongoose [12]. The contact-era archaeological site En Bas Saline (Haiti) contains the first Caribbean occurrence of the black rat (*Rattus rattus) *[13]. Isolated jaguar teeth found at Punta Candelero (Late Ceramic), Puerto Rico, suggest post-mortem trade from South America [14]. Thirty rhesus macaques* (Macaca mulatta) *were imported to Cayo Santiago in 1938 for medical studies, expanding to >1000 animals today [15]*. *D. In 1995, 15 green iguanas (*Iguana iguana, E. LA*) rafted from Guadeloupe to Anguilla during a hurricane [16]; this event was contingent on the past introduction of green iguanas by Indigenous and/or colonial groups [17]. The Critically Endangered Antiguan racer (*Alsophis antiguae) *has been translocated to three offshore islets, growing the population from 51 to >1100 individuals [18]. In St. Kitts, colonial French brought African green monkeys (*Chlorocebus aethiops sabaeus, E. LA) *as pets in ~1670; troops persist today [19]. Isotopic data suggest colonial Spanish/Portuguese translocated peccaries (Tayassuidae) from the Guianas to Barbados post-1600 [20]. Specimens from the Early Ceramic site Pearls, Grenada, suggest the possible translocation of small rice rats* (Zygodontomys sp.) *from South America [21]*.

Here, we synthesize paleontological, archeological, and historical data from the Caribbean to provide the first continuous dataset of vertebrate species introductions spanning ~7,000 years ago into the present day, thus providing a moving image of the formation of an insular Anthropocene biota from initial human arrival. While other studies have looked at the past or present in isolation, we show that Indigenous people had distinct impacts on their local ecosystems across different islands and cultural time periods that laid the foundation for subsequent species introductions mediated by Europeans and ongoing introductions through industrial trade. We quantified the timing, spread, and diversity of non-native terrestrial vertebrates throughout the Caribbean as they relate to cultural practices and identify prospects for conservation across the heterogeneous present-day landscape.

### Navigating interdisciplinary datasets and terminologies

We recognize there is a broad body of literature that classifies species introductions by origin and stage of the invasion process, but for the purposes of this paper, we define introduced species as species that have been translocated to an island where a native, extant population of the species is not currently present [22]. We consider both the introduction of species to islands that they have never been native to as well as re-introductions of native species that had been extirpated previously. We refer to non-native species collectively as “introduced species,” “non-native species,” and “translocated species” throughout this paper, because our primary interest is in documenting the accumulation of species through human activities, and only use the term “invasive species” when the species is recognized as such by an outside authority.

We compiled a database of terrestrial vertebrate species that were introduced to the Caribbean ([23–27]; Supplementary Tables 1 - 4) that records the species name, place of origin, and the location of the introduced populations (both the specific island and its place within the Bahamas, Greater Antilles, or Lesser Antilles). Due to ambiguity associated with the exact locality of origin, we used broad biogeographic realms (e.g., Neotropics). Where possible, we included the introduction date, number of times a species was introduced to a particular locality, reason for introduction, and whether an introduction was successful. We discuss three distinct periods that loosely correspond to different societies and economic structures: Indigenous societies, colonial societies, and modern societies, and identify relevant cultural and geographic groupings within these periods.

While we are able to present modern and colonial introductions at the centennial scale, the coarser data resolution for introductions by Indigenous groups required that we consider these data as one-time bin for quantitative analyses. We acknowledge the diversity present in the Caribbean through time and discuss these patterns qualitatively using archaeological groupings rather than particular ethonyms, as there is no clear consensus regarding the linguistic, genetic, and cultural origins of many groups. Archaeologists typically recognize four temporal bins when discussing waves of Caribbean colonization and compare groups based on these time periods, rather than cultural names: Lithic (~5-7,000 years ago), Archaic (~2-5,000 years ago), Ceramic (~2,500 years ago in Puerto Rico and the Virgin Islands, with later dates elsewhere), and lastly, the historical period starting with the arrival of Columbus in 1492 AD [28].

The first people arrived in the Caribbean ~7,000 years ago, via the South American Orinoco Basin and/or Central America [28]. The earliest Lithics sites found in the Greater Antilles (Cuba and Hispaniola), though poorly studied, are marked by a flaked-stone technology, and are largely lithic workshops that lack human skeletal remains [28]. Subsequently, Archaic populations arrived from northeastern South America, and represented both marine and terrestrial subsistence strategies [29]. Increasing evidence supports food management if not outright cultivation, including digging stone tools and a variety of plants transported outside of their natural range, as well as large area burning and early evidence of pottery [30].

The Ceramic Age – a time of increased sedentism and complex trade networks – is demarcated ~2000 years ago by the arrival of “Saladoid” groups from South America into the Lesser Antilles and Puerto Rico [28]. These Arawak-speaking peoples from the Orinoco Valley may have admixed with, rather than replaced, existing Lithic/Archaic populations [30]. By the 15^th^ century, Europeans estimated indigenous population sizes of 100,000 to 6 million [31]. While Europeans have labeled many of these groups upon contact with various ethnonyms, such as Taino (Greater Antilles), Lucayan (Bahamas), and Caribs (Lesser Antilles, coastal South America), these labels were often problematic in their misinterpretation of existing political, ethnic, and linguistic diversity [32].

Colonial societies span 1492 to 1800 and represent a time of massive demographic turnover that accompanied the onset of large-scale colonial agriculture , with forced relocation of indigenous people across the Antilles, temporary depopulation of the Bahamas, disease, slavery, and war, leading to indigenous population decreases and admixture with other American populations, as well as European and enslaved African populations (though Indigenous genetic lineages persist today [33]). The Spanish arrived first but they were soon followed by Dutch, British, and French colonizers, all of whom formed complex trade networks within the Caribbean and across the Atlantic Ocean, further connecting the archipelago to continental North America, Europe, and Africa. The mass influx of people from colonization, enforcement of the encomienda system, and the transatlantic slave trade produced social and economic stratifications still seen today.

We use the midpoint of the Industrial Revolution, 1800, to demarcate the onset of modern society, as innovations in technology and machinery drastically changed worldwide economies and the connectivity of regions. This time period roughly corresponds to a number of significant events unfolding in the Caribbean: the Haitian revolution (1791 - 1804) which led to slavery abolition and Haitian independence, and the Slave Trade act of 1807, which prohibited the trade of enslaved people in the British Empire. The abolition of slavery spurred the recruitment and migration of people from Asia as indentured servants, most notably China and India. The majority of Caribbean nations did not achieve independence until the 20th century , and select islands remain overseas territories today.

### Species accumulation over space and time

Species introductions in the Caribbean accelerate towards the present in all terrestrial vertebrate groups, but the timing of these changes vary across taxa (Figure 2). Whereas mammal introductions start during the Indigenous period and continue to increase throughout the colonial and modern period, confirmed bird introductions only begin during the colonial period but quickly surpass all other groups. There are several pre-modern reptile introductions, but the majority of introduced reptiles arrive during the modern period, as do amphibians. As the number of introduced species increases, so does the range of biogeographic origins represented (Figure 3). Species introductions by Indigneous groups are exclusively Neotropical or Nearctic in origin, and by the 21st century taxa from every temperate and tropical biogeographical realm have been introduced into every Caribbean sub-archipelago.

**Figure 2.**
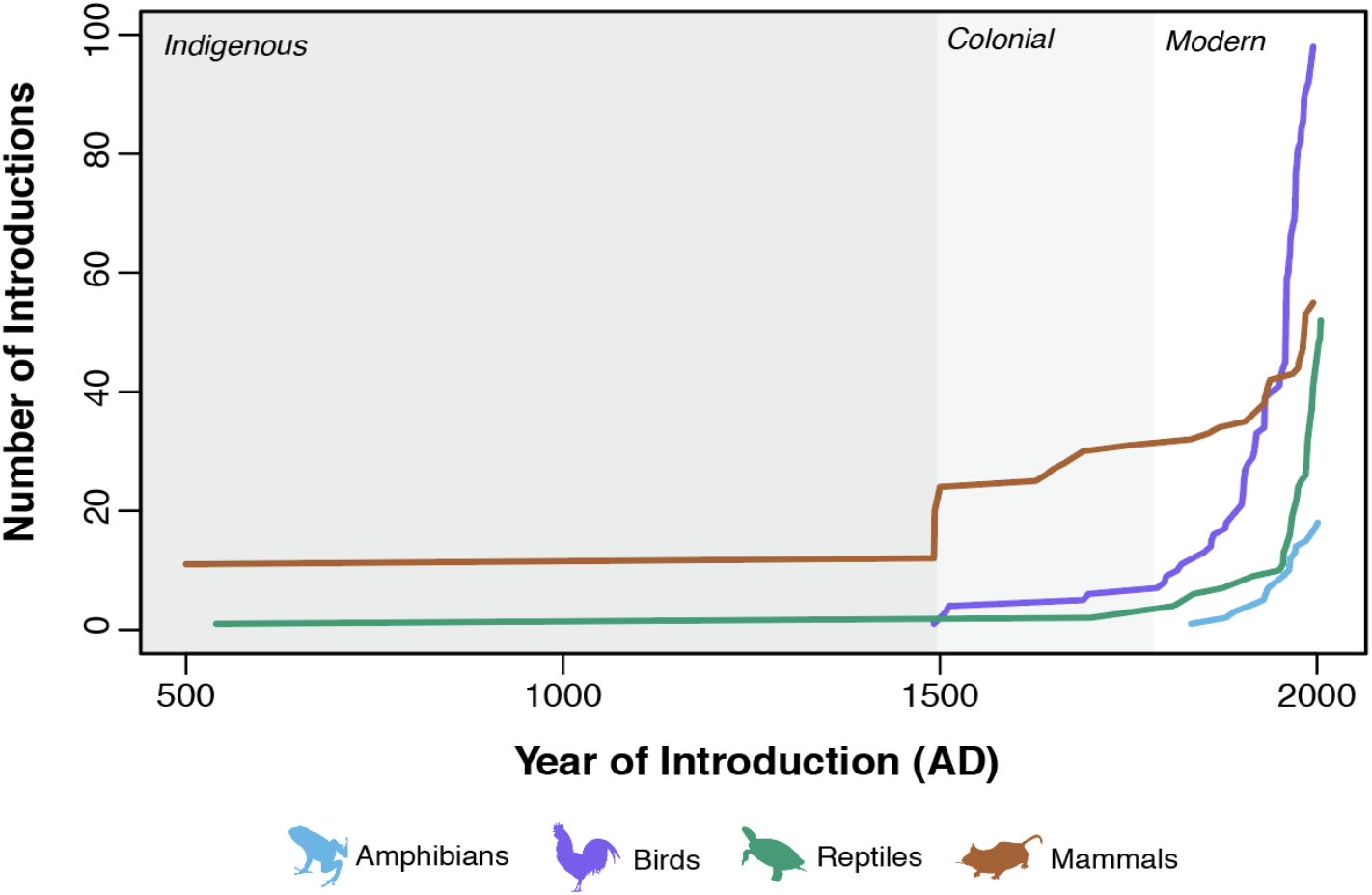
Accumulation of introduced species over time, beginning with the first dated introduction for each species in our database

**Figure 3.**
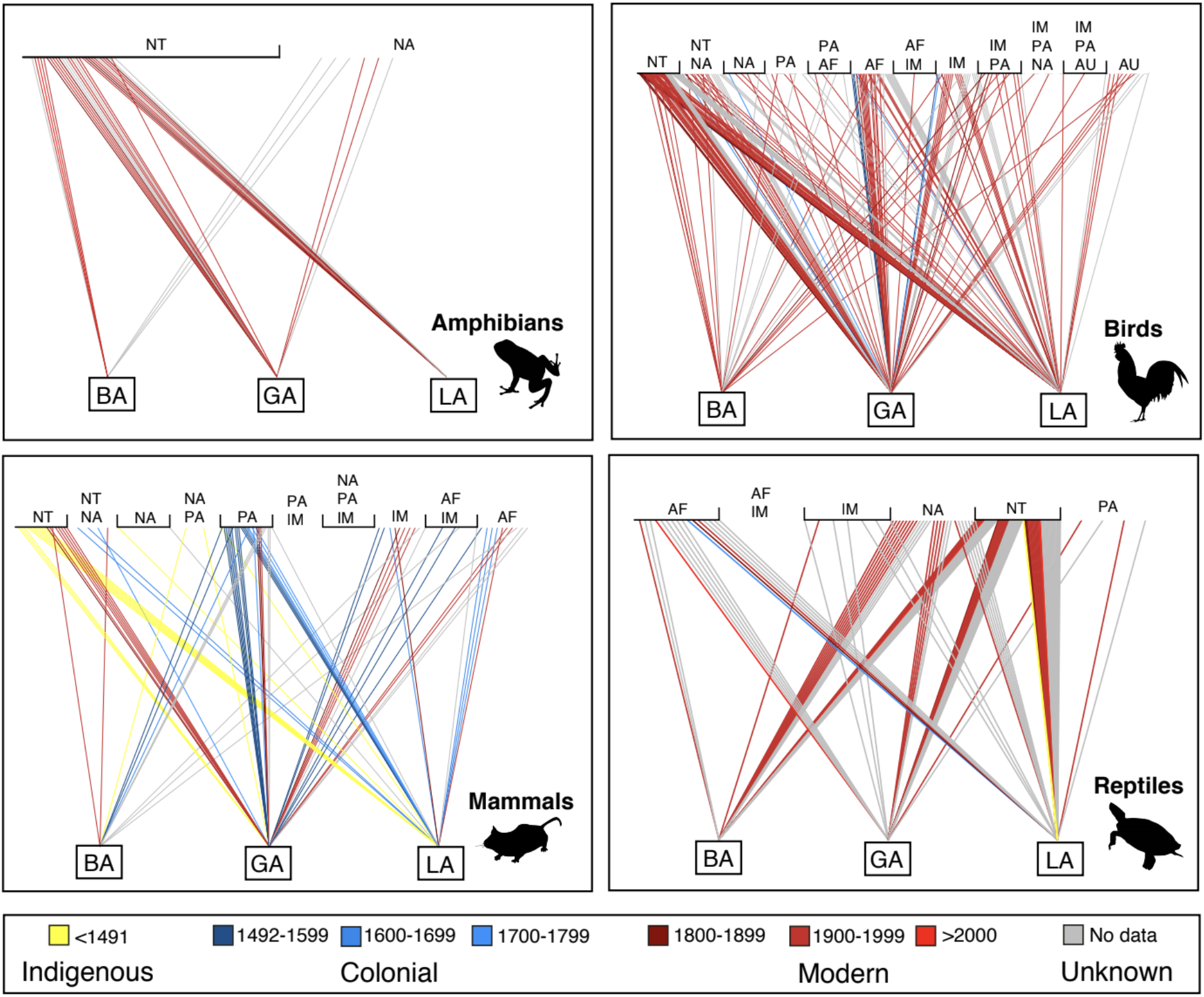
Bipartite network plots showing the biogeographic origin of species introduced to the Bahamas (BA), Greater Antilles (GA), and Lesser Antilles (LA), color coded by time of introduction. Abbreviations: NT = Neotropical, NA = Nearctic, PA = Palarctic, IM = Indo-Malay, AF = Afrotropical, AU = Australasian.

### Ambiguity in introduction chronology and species origin

It is currently impossible to reconstruct a fully dated introduction chronology given the high level of uncertainty in the Caribbean record (due to a lack of radiocarbon dates, genetic estimates, historic documentation, etc). Mammals have the best coverage, with 88% of taxa having a first introduction date. This contrasts with reptiles, amphibians, birds, and reptiles, where only 69%, 66%, and 49% of first introductions have a date, respectively. This ambiguity in the timing of a given occurrence also propagates uncertainty when delimiting whether a species is in fact native or not, as it may be unclear whether a species appears before, concurrent with, or after the arrival of humans or a given culture on an island.

We identify introduced species based on an understanding of a species’ native biogeographic range. Delimiting the native biogeographic range, however, requires a thorough understanding of pre-human paleontological baselines as ranges shifts over time and can be cryptically extended by undocumented human activities. For example, the number of iguana species, *Iguana* sp., native to the Lesser Antilles have been a source of contention in part because of ambiguity in biogeography. Two species in this genus are presently found in the Caribbean: *Iguana delicatissima*, which is considered endemic to the Lesser Antilles, and the green iguana, *Iguana iguana*, which is widely distributed throughout the Caribbean and Central America. *Iguana iguana* is considered an invasive species and can hybridize with endemic iguanas including both *I. delicatissima* [34] and rock iguanas of the genus *Cyclura* [35]. Until recently, *I. iguana* was believed to have been deliberately introduced in historical times [36]. However, a reanalysis of zooarchaeological materials from Guadeloupe indicate a pre-Columbian introduction of *I. iguana* from Central America in at least this part of the Lesser Antilles [17], though it is not clear whether there is a direct ancestor-descendent relationship between these remains and extant populations, or if there have been subsequent colonizations.

When a specimen of a putatively non-native species is found in an archaeological context, there is still uncertainty as to whether the translocation represents the movement of a live or dead animal, as animal parts were often traded post-mortem as materials; this dichotomy has clear consequences for ecological impacts and population establishment. For example, [37] showed that modified deer (*Odocoileus*, *Mazama*) bones from 29 Late Ceramic sites in the Lesser Antilles represent ornamental material rather than the importation of living deer. Where deer are native on Trinidad, their skeletal elements are found in abundance and burned in archaeological assemblages; conversely, where they are not native, the diversity of skeletal elements represented is low, the elements present are relatively rare in the assemblages, and they are modified, rather than burned.

The trading networks of Indigenous peoples have shaped patterns of standing biodiversity worldwide [22], and if not accounted for, can obfuscate the true ecological and evolutionary processes underlying the generation of diversity and miss an opportunity to understand how humans created communities prior to European arrival. While *Iguana delicatissima* is assumed to be endemic to the Lesser Antilles as a whole (as compared with isolated islands), it is only known from cultural sites in the paleontological record. The Kalinago/Carib name for St. Lucia (Hewanorra) translates to *Land of the Iguanas*, suggesting that they encountered higher abundances of iguanas there than on other islands between South America and St. Lucia [38]. It is unclear whether Indigenous populations then translocated *Iguana delicatissima* as they moved throughout the Lesser Antilles, and if so, which island represents the original source population. Similar intra-Caribbean translocations occurred for endemic rodents of the Greater Antilles (Figure 1 [39]).

Investigating the native status of certain species is an active area of research, and in numerous cases, we were unable to include examples in our dataset due to lack of suitable paleontological baselines or zooarchaeological evidence. For example, the critically endangered “mountain chicken” (a large frog, *Leptodactylus fallax*), a prized food source restricted today to Dominica and Montserrat, may have been translocated by pre-Columbian to St. Vincent, Martinique, and Grenada [38], but more evidence is necessary to corroborate this claim.

### Human niche construction and historically contingent outcomes in the Caribbean

Species introductions represent a key component of human niche construction, influencing the behavior, ecology, and evolution of humans and other species [5]. Because each round of human colonization of the Caribbean results in new niche construction activities, the long-term effects compound and form a new ecological inheritance for subsequent human colonizers and ecological communities [40]. Historical contingency—the effect of the order and timing of species introductions—influences community assembly processes, and priority effects allow early arriving species to fill or alter niches and thereby discourage the integration of additional species [41]. In this logic, the presence of native fauna may delay the establishment of introduced species that occupy similar niches; therefore, pre-European extinctions of native fauna may have inadvertently provided an opening for colonial introduced species to become established. Below we briefly review some key examples of niche construction across our three temporal groupings.

#### Indigenous Societies (pre-1492)

Indigenous societies introduced species for direct consumption and material use. All of these introduced species/lineages are of Neotropical origins, which makes distinguishing them from native taxa difficult in the absence of pre-human baselines. The number of introduced species contributed by Indigenous people may thus be an underestimation, especially because many of these taxa are now extirpated in their introduced range, and our knowledge of these introductions is restricted to a potentially undersampled archaeological record.

Vertebrate translocations began in the Archaic: the Puerto Rican hutia, †*Isolobodon portoricensis*, is mistakenly named, as it was cryptically introduced from Hispaniola into Puerto Rico as early as >3000 years ago, followed by a subsequent introduction to St. John (Virgin Islands) several hundred years later [42],[43]. Isotopic and radiocarbon evidence also corroborates the multiple intra-Bahamas translocations and Ceramic age use of the Bahamian hutia, *Geocapromys ingrahami* [39], which may have contributed to its persistence into the present day by creating metapopulations to buffer the effects of overhunting and hurricanes (Figure 1).

The guinea pig (*Cavia porcellus*) was introduced to the Caribbean during the Late Ceramic age, and is known from 18 sites across 10 islands, with an earliest appearance in Puerto Rico more than a thousand years ago [44]. Their low abundance in middens suggests they were used as a supplemental, not primary, food source, and mitogenomic data indicate multiple introductions occurred from South America, [44]. Dogs became widespread in the Caribbean during the Ceramic period, and strontium isotopes detected the presence of both local and nonlocal dogs and humans at Anse a la Gourde and Morel, two sites on Guadeloupe [45]. Together, these examples confirm continued movement and trade between South America and the Caribbean at this time.

As many of these translocated species have not persisted into the present day, the impacts of these ephemeral introductions on past endemic species remains poorly understood. Congruent with global patterns of insular extinction, a number of endemic species go extinct shortly after the arrival of people in the archipelago [10]. Thus, extinction and habitat modification may have primed insular systems like the Caribbean for subsequent species introductions and establishment.

#### Colonial Societies (1492 - 1799)

Colonial societies introduced a number of taxa for food, often releasing populations of animals such as goats and pigs that then rapidly established feral populations. There were also inadvertent introductions of human commensals such as the black rat (*Rattus rattus*) and house mouse (*Mus musculus*). Agricultural introductions and practices during this time period heavily altered native vegetation [46, 47]. Sugarcane, native to southeast Asia, was introduced throughout the Caribbean during the 16th and 17th centuries. Towards the later years of the colonial era, Caribbean economies began to implement intensive monocrop sugarcane agriculture to keep up with demand, initiating the trend towards widespread deforestation. Tobacco (*Nicotiana tabacum*), which was first introduced to the Caribbean by Indigenous people during the Ceramic Age, also became an increasingly important crop during this time for European markets, particularly in the Dominican Republic and Puerto Rico [42],[43, 47].

There was also an explicit desire to design new ecosystems, as exemplified by naturalists like Félix Louis L’Herminier. L’Herminier and his son collected fauna from Puerto Rico and West Africa and intentionally introduced them to Guadeloupe in acclimatization experiments [38]. Colonial societies were also introducing taxa from abroad to make their new environment more familiar, meet aesthetic expectations, and support leisure practices such as hunting and pet keeping [23, 24]. At this point we begin to see the breakdown of biogeographic realms, as the majority of introduced vertebrates come from the Palearctic, Afrotropics, and Indo-Malay regions rather than the Neotropics (Figure 3).

#### Modern Societies (1800 - present)

Crops cultivated by colonial societies continued to play a significant role in the Caribbean decades later. By 1900, eleven Caribbean territories across the Greater and Lesser Antilles were dependent on sugarcane production [48], in turn generating an economic impetus for pest control. Mongooses (*Herpestes auropunctatus*) were introduced to control the rat populations (*Rattus* sp.) that began with contact in 1492 (Figure 1) and cane toads (*Rhinella marina*) were introduced to control both native and introduced insect pests; both species were largely ineffective as pest control agents. Instead, they contributed to the extinction of endemic vertebrate lineages across the region and continue to pose significant conservation challenges today [49, 50].

Reptile and amphibian introductions have become more commonplace in the past century as a result of largely accidental events facilitated by trade [3]. When purposefully translocated, the intent is usually that they remain contained (e.g. pet trade), though there have been escapes. Following scientific awareness of the detrimental effects of invasive species, most intentional introductions during this time period are for a conservation goal, such as re-establishing an extirpated species [18], although there are numerous occasions in which taxa are introduced onto small islands and cays for basic scientific research [51]. Moving into the present day, we see the complete breakdown of biogeographic barriers, the wide establishment of cosmopolitan commensals, landscapes dominated by agriculture and livestock, and the formation of an Anthropocene fauna.

### Present day biodiversity: a patchwork of past movements and practices

Centuries of human movement and species translocations in the Caribbean have resulted in a biological patchwork of novel ecological communities, reflecting the heterogeneous distribution of economic and cultural activities throughout the region and through time. Our analyses reveal introduced species from all corners of the world on islands of every size, substrate, and age; few ecological communities have been unmarred by species introductions, and the islands without introduced species tend to be small, low-lying, islands uninhabited by humans that face frequent disturbances from hurricane activity. The patchy distributions of some groups throughout the Caribbean, such as ground-dwelling lizards and snakes, may have resulted from different biocontrol regimes instituted by different societies, as well as differential usages of islands. For example, the usage of La Desirade (Lesser Antilles) as a leper colony as opposed to sugarcane production may have facilitated the survival of *Iguana delicatissima* on the island, even though it has been extirpated from nearby Guadeloupe, where sugar plantations were common. In some places, the contrast of different human cultural groups and economic practices can be seen across the same landscape. Hispaniola is currently divided into two nations with approximately the same population size but differing areas: the larger Dominican Republic in the east and smaller Haiti in the west. Hispaniola’s geotectonic history naturally split species into north/south rather than east/west ranges, resulting in species that are distributed across similar ecosystems but face unequal anthropogenic threats built on centuries of differences across a colonial boundary [52].

### Introductions, extinctions, and the restoration of ecosystem functions

Paleontological studies have identified patterns of size-biased vertebrate extinction in the Caribbean [53],[10], although we are still elucidating extinction chronologies for many taxonomic groups. Large-bodied terrestrial mammals such as ground sloths disappeared prior to the Ceramic period, potentially linked to hunting or habitat alteration [10]. The introduction of species during the Indigenous period does not appear to have caused extinction directly. Conversely, the size-biased loss of small-bodied mammals across the region, including the eulipotyphlan “island shrews” (†Nesophontidae) and a diverse group of rodent species including hutias (Capromyidae), spiny rats (Echimyidae), and rice rats (Cricetidae) corresponds well to the arrival of non-native mammalian carnivores and *Rattus*. This introduction of potential predators and competitors links both accidental and intentional species introductions to a major extinction event in the Caribbean mammals. Similarly, the disappearance of many lizards occurred after European arrival [53],[49], likely due to impacts from introduced species.

Species reintroductions and eradications are currently being conducted in an effort to restore altered ecosystems to a previous natural state, but our data highlighting millennia of introductions beg the question: what previous state is the “natural” state? At what cultural grouping or introduction do Caribbean ecosystems move from being “natural” to “novel”, and what defines a suitable restoration target? From a deeper time perspective, it is clear that we can never return to a pre-human or pre-European state in the Caribbean given the disappearance of many unique evolutionary lineages during the Holocene, though it may be possible to replace the ecological function of some of these lost species. Endemic ground sloths and monkeys may have shaped vegetation structure through browsing and potential mutualistic relationships; their extinction may have opened space for non-native large mammals, especially small ungulates and the non-native primates that have now established colonies on some of the islands (Figure 1).

Many prehistoric extinctions have been linked to a loss in ecosystem services [54]. Frugivores provide many ecosystem services related to pollination and seed dispersal, and a number of Caribbean frugivores, including bats, birds, sloths, and tortoises, went extinct both before and after European arrival [10]. Extinctions in these groups may also have impacted vegetation structure, insect diversity, and insect abundance. Recent research in other insular systems suggests that introduced birds can serve as important seed dispersers for native plants that may have lost their original dispersal agents [55], a relevant finding for Caribbean plants given the loss of native seed dispersers and arrival of introduced birds.

Many Caribbean landscapes meet the definition of a novel ecosystem in being dominated by anthropogenic alterations [56]. In isolated areas such as national parks and/or difficult terrain, remnants of historic ecosystems may persist, though both the ecosystems themselves and our understanding of their “natural” state continue to change with accelerating human activity, increased exploration of paleontological and archaeological records, and anthropogenic climate change. If the goal is to maintain the status quo of insular biodiversity and protect still-surviving endemic species, eradication of non-native species represents the most significant gain for island endemics [57]. A global analysis of insular systems indicated that the Caribbean has a large proportion of biologically important islands (45 of 292, or 15.4%) where eradication would be beneficial [58]. Ongoing eradications of *Rattus* from small islands used heavily by seabirds may have positive ecological effects on nutrient cycling [59]. Even when an island-wide eradication is unfeasible, local actions can have large conservation impacts; a dog cull in the Duchity region of Haiti has benefited Hispaniolan solenodons [60].

In addition to removing introduced species, conservation efforts are re-introducing recently extirpated species or introducing closely related species. Several reintroductions of extirpated species have been successful [18], but these endeavors typically necessitate eradications of predators and changes to law enforcement. While conservation programs have brought the Grand Cayman blue iguana (*Cyclura lewisi*) and Puerto Rican parrot (*Amazona vitatta*) back from the brink of extinction, they are again victims of cats, dogs, humans, and cars once outside of protected release areas [61],[62]. These conservation efforts beg the question of if and how reintroductions carried out today recapitulate the intra-Caribbean movement of species by Indigenous groups thousands of years ago – some of these translocated populations like the Bahamian hutia have persisted, whereas others, like the Puerto Rican hutia, have not. The lessons from these past translocations have yet to be fully leveraged in regional conservation decision-making processes or prioritization schemes.

## Summary

Humans significantly altered the Caribbean’s biodiversity since first arriving in the archipelago, homogenizing its once evolutionarily isolated, endemic fauna with other biogeographic realms. The introduction chronology presented here across three broad categories—Indigenous, colonial, and modern— compliments known extinction chronologies for Caribbean taxa, and together these data provide a glimpse into shifting community structure, species interactions, and ecosystem function. We find that the present-day Caribbean fauna is historically contingent on this chronology of human occupation and the diverse cultural practices of these different groups. While introductions during the colonial and modern period have been previously linked to extinction events in the native fauna, attention is shifting towards understanding the ecological impacts of prehistoric extinctions, and how these extinctions may have laid the groundwork for subsequent introduction events. Introductions begot more introductions both directly and indirectly, such as the accidental introduction of a pest that required an intentionally introduced predator to control it.

Invasive species represent one of the greatest management challenges facing insular biota globally [57, 58]. Our synthesis builds on a body of literature showing that the presence of non-native species in the context of altered landscapes and growing human populations is not a new phenomenon; rather, the predisposition of certain insular ecosystems to the effects of modern invasive species may be undergirded by their unique accumulated histories of Indigenous and colonial activity, demanding inclusion of paleontological and archaeological knowledge in conservation planning [56]. What lessons can we learn about coexistence with non-native species from this past record of human niche construction? Such line of thought is particularly vital given that invasive species removals can have unintended ecological consequences if the whole system is not taken into account [63].

It is important to note that the archaeological record shows substantial variation across Indigenous communities over space and time, which our analyses were not able to address due to the current resolution of the zooarchaeological record, such as differences in human-environment interactions among early arriving Lithic/Archaic groups. Deepening our understanding of translocations and in particular, understanding multiple introductions or demographic trends of relevance to ecological hypotheses, requires increased resolution in the radiocarbon record and a combination of multiple lines of evidence, including ancient DNA, collagen sequencing, isotopes, and dental calculus from the archaeological and paleontological records [64]. While we have focused on terrestrial vertebrates, the abundance of shell middens throughout the Caribbean present a clear opportunity to provide context for ongoing marine species invasions.

More generally, because of taphonomic processes and biases in written history, the absence of evidence for certain intensive practices cannot be used to argue that Indigenous populations did not dramatically shape their environments. This reality often conflicts with the widespread usage of 1492 as a restoration baseline in North American conservation practices, which ignores the impacts of Indigenous people who were modifying landscapes long before European arrival. Paleontological and zooarchaeological datasets provide tools for reconstructing pre-human environments and evaluating the consequences of anthropogenic modifications and species introductions over the long-term. Knowing that present-day communities are historically contingent tells us why some places retain high levels of native biodiversity whereas other areas are dominated by non-native fauna, and can help us pinpoint conservation opportunities in a rapidly changing and increasingly connected world.

## Acknowledgements

We thank the numerous archaeologists whose collections made our analyses possible. K Rayfield, R Singleton, K Hughes, and R Austin provided helpful comments that enriched the text of this manuscript. Image credits: Phylopic; hutia by Mark Erdos/IUCN SMSG; green monkey by Rod Waddington; green iguana by Alan Schmierer; northern bobwhite by Steve Maslowski/USFWS.

## References

[1] Erlandson, J.M. & Braje, T.J. 2013 Archeology and the Anthropocene. Anthropocene 4, 1–7. (doi:10.1016/j.ancene.2014.05.003).

[2] Hobbs, R.J., Higgs, E. & Harris, J.A. 2009 Novel ecosystems: implications for conservation and restoration. Trends in ecology & evolution 24, 599–605. (doi:10.1016/j.tree.2009.05.012).

[3] Helmus, M.R., Mahler, D.L. & Losos, J.B. 2014 Island biogeography of the Anthropocene. Nature 513, 543–+. (doi:10.1038/nature13739).

[4] Anderson, A. 2009 The rat and the octopus: initial human colonization and the prehistoric introduction of domestic animals to Remote Oceania. Biological Invasions 11, 1503–1519. (doi:10.1007/s10530-008-9403-2).

[5] Boivin, N.L., Zeder, M.A., Fuller, D.Q., Crowther, A., Larson, G., Erlandson, J.M., Denham, T. & Petraglia, M.D. 2016 Ecological consequences of human niche construction: Examining long-term anthropogenic shaping of global species distributions. Proceedings of the National Academy of Sciences of the United States of America 113, 6388–6396. (doi:10.1073/pnas.1525200113).

[6] Hulme, P.E. 2009 Trade, transport and trouble: managing invasive species pathways in an era of globalization. Journal of Applied Ecology 46, 10–18. (doi:10.1111/j.1365-2664.2008.01600.x).

[7] Stuble, K.L. & Souza, L. 2016 Priority effects: natives, but not exotics, pay to arrive late. Journal of Ecology 104, 987–993. (doi:10.1111/1365-2745.12583).

[8] Turvey, S.T. & Saupe, E.E. 2019 Insights from the past: unique opportunity or foreign country? Philosophical Transactions of the Royal Society B-Biological Sciences 374, 20190208. (doi:10.1098/rstb.2019.0208).

[9] Myers, N., Mittermeier, R.A., Mittermeier, C.G., da Fonseca, G.A.B. & Kent, J. 2000 Biodiversity hotspots for conservation priorities. Nature 403, 853–858.

[10] Cooke, S.B., Davalos, L.M., Mychajliw, A.M., Turvey, S.T. & Upham, N.S. 2017 Anthropogenic Extinction Dominates Holocene Declines of West Indian Mammals. In Annual Review of Ecology, Evolution, and Systematics, Vol 48 (ed. D.J. Futuyma), pp. 301–327.

[11] LeFebvre, M.J., DuChemin, G., deFrance, S.D., Keegan, W.F. & Walczesky, K. 2019 Bahamian hutia (Geocapromys ingrahami) in the Lucayan Realm: Pre-Columbian Exploitation and Translocation. Environmental Archaeology 24, 115–131. (doi:10.1080/14614103.2018.1503809).

[12] Scott, W. 1982 Observations on the birds of Jamaica, West Indies. The Auk 9, 273–277.

[13] Deagan, K. 2004 Reconsidering Taino social dynamics after Spanish conquest: Gender and class in culture contact studies. American Antiquity 69, 597–626. (doi:10.2307/4128440).

[14] Laffoon, J.E., Rodríguez Ramos, R., Chanlatte Baik, L., Narganes Storde, Y., Rodríguez Lopez, M., Davies, G.R. & Hofman, C.L. 2014 Long-distance exchange in the precolonial Circum-Caribbean: A multi-isotope study of animal tooth pendants from Puerto Rico. Journal of Anthropological Archaeology 35, 220–233. (doi:10.1016/j.jaa.2014.06.004).

[15] Rawlins, R.G. & Kessler, M.J. 1986 The Cayo Santiago macaques: History, behavior, and biology, SUnY Press.

[16] Censky, E., Hodge, K. & Dudley, J. 1998 Over-water dispersal of lizards due to hurricanes. Nature 395, 556. (doi:10.1038/26886).

[17] Bochaton, C., Bailon, S., Ineich, I., Breuil, M., Tresset, A. & Grouard, S. 2016 From a thriving past to an uncertain future: Zooarchaeological evidence of two millennia of human impact on a large emblematic lizard (Iguana delicatissima) on the Guadeloupe Islands (French West Indies). Quaternary Science Reviews 150, 172–183. (doi:10.1016/j.quascirev.2016.08.017).

[18] Daltry, J.C., Lindsay, K., Lawrence, S.N., Morton, M.N., Otto, A. & Thibou, A. 2017 Successful reintroduction of the Critically Endangered Antiguan racer Alsophis antiguae to offshore islands in Antigua, West Indies.(Report). International Zoo Yearbook 51, 97. (doi:10.1111/izy.12153).

[19] McGuire, M.T. 2017 The St. Kitts Vervet (Cercopithecus aethiops). Journal of Medical Primatology 3, 285–297. (doi:10.1159/000460030).

[20] Giovas, C.M., Kamenov, G.D. & Krigbaum, J. 2019 87Sr/86Sr and 14C evidence for peccary (Tayassuidae) introduction challenges accepted historical interpretation of the 1657 Ligon map of Barbados. PloS one 14, e0216458–e0216458. (doi:10.1371/journal.pone.0216458).

[21] Mistretta, B.A. 2019 Grenada’s extinct rice rats (Oryzomyini): Zooarchaeological evidence for taxonomic diversity. Journal of Archaeological Science: Reports 24, 71–79. (doi:10.1016/j.jasrep.2018.12.018).

[22] Hofman, C. & Rick, T. 2018 Ancient Biological Invasions and Island Ecosystems: Tracking Translocations of Wild Plants and Animals. Journal of Archaeological Research 26, 65–115. (doi:10.1007/s10814-017-9105-3).

[23] Long, J.L. 1981 Introduced birds of the world: the worldwide history, distribution and influence of birds introduced to new environments / by John L. Long; illustrated by Susan Tingay. New York, Universe Books.

[24] Long, J.L. 2003 Introduced mammals of the world: their history, distribution and influence / John L. Long. Collingwood, Vic., Australia, CSIRO Pub.

[25] Kraus, F. 2009 Alien Reptiles and Amphibians a Scientific Compendium and Analysis / by Fred Kraus. 1st ed. 2009. ed. Dordrecht, Springer Netherlands.

[26] Powell, R., Henderson, R.W., Farmer, M.C., Breuil, M., Echternacht, A.C., van Buurt, G., Romagosa, C.M. & Perry, G. 2011 Introduced amphibians and reptiles in the greater Caribbean: Patterns and conservation implications63–143 p.

[27] Dyer, E.E., Redding, D.W. & Blackburn, T.M. 2017 The global avian invasions atlas, a database of alien bird distributions worldwide. Scientific Data 4, 170041. (doi:10.1038/sdata.2017.41).

[28] Keegan, W.F.H., Corinne L. 2017 The Caribbean before Columbus. New York, NY, Oxford University Press.

[29] Reid, B.A. 2018 The Archaeology of Caribbean and Circum-Caribbean Farmers (6000 BC - AD 1500), Taylor and Francis.

[30] Keegan, W.F. 2006 Archaic influences in the origins and development of Taino societies. Caribbean Journal of Science 42, 1–10.

[31] Keegan, W. 1996 West Indian archaeology. 2. After Columbus. Journal of Archaeological Research 4, 265–294. (doi:10.1007/BF02229089).

[32] Hofman, C.L., Hung, J.U., Malatesta, E.H., Jean, J.S., Sonnemann, T. & Hoogland, M. 2018 Indigenous Caribbean perspectives: archaeologies and legacies of the first colonised region in the New World.(Report). Antiquity 92, 200. (doi:10.15184/aqy.2017.247).

[33] Nieves-Colón, M.A., Pestle, W.J., Reynolds, A.W., Llamas, B., de La Fuente, C., Fowler, K., Skerry, K.M., Crespo-Torres, E., Bustamante, C.D. & Stone, A.C. 2019 Ancient DNA reconstructs the genetic legacies of pre-contact Puerto Rico communities. Molecular biology and evolution. (doi:10.1093/molbev/msz267).

[34] Vuillaume, B., Valette, V., Lepais, O., Grandjean, F. & Breuil, M. 2015 Genetic Evidence of Hybridization between the Endangered Native Species Iguana delicatissima and the Invasive Iguana iguana (Reptilia, Iguanidae) in the Lesser Antilles: Management Implications. Plos One 10, e0127575. (doi:10.1371/journal.pone.0127575).

[35] Moss, J.B., Welch, M.E., Burton, F.J., Vallee, M.V., Houlcroft, E.W., Laaser, T. & Gerber, G.P. 2018 First evidence for crossbreeding between invasive Iguana iguana and the native rock iguana (Genus Cyclura) on Little Cayman Island. Biological Invasions 20, 817–823. (doi:10.1007/s10530-017-1602-2).

[36] Stephen, C.L., Reynoso, V.H., Collett, W.S., Hasbun, C.R. & Breinholt, J.W. 2013 Geographical structure and cryptic lineages within common green iguanas, Iguana iguana. Journal of Biogeography 40, 50–62. (doi:10.1111/j.1365-2699.2012.02780.x).

[37] Giovas, C.M. 2018 CONTINENTAL CONNECTIONS AND INSULAR DISTRIBUTIONS: DEER BONE ARTIFACTS OF THE PRECOLUMBIAN WEST INDIES - A REVIEW AND SYNTHESIS WITH NEW RECORDS.(Report). Latin American Antiquity 29, 27. (doi:10.1017/laq.2017.57).

[38] Breuil, M. 2002 Histoire Naturelle des Amphibiens et Reptiles Terrestres de l’Archipel Guadeloupéen: Guadeloupe, Saint-Martin, Saint-Barthélemy. Paris, Muséum National d’Histoire Naturelle.

[39] LeFebvre, M.J., deFrance, S.D., Kamenov, G.D., Keegan, W.F. & Krigbaum, J. 2019 The zooarchaeology and isotopic ecology of the Bahamian hutia (Geocapromys ingrahami): Evidence for pre-Columbian anthropogenic management.(Research Article). PLoS ONE 14, e0220284. (doi:10.1371/journal.pone.0220284).

[40] Kendal, J., Tehrani, J.J. & Odling-Smee, J. 2011 Human niche construction in interdisciplinary focus Introduction. Philosophical Transactions of the Royal Society B-Biological Sciences 366, 785–792. (doi:10.1098/rstb.2010.0306).

[41] Fukami, T. 2015 Historical Contingency in Community Assembly: Integrating Niches, Species Pools, and Priority Effects. In Annual Review of Ecology, Evolution, and Systematics, Vol 46 (ed. D.J. Futuyma), pp. 1–23.

[42] Newsom, L.A. & Wing, E.S. 2004 On land and sea: Native American uses of biological resources in the West Indies. Tuscaloosa, University of Alabama Press.

[43] Rivera-Collazo, I.C. 2015 Por el camino verde: Long-term tropical socioecosystem dynamics and the Anthropocene as seen from Puerto Rico. Holocene 25, 1604–1611. (doi:10.1177/0959683615588373).

[44] Lord, E., Collins, C., deFrance, S., LeFebvre, M.J. & Matisoo-Smith, E. 2018 Complete mitogenomes of ancient Caribbean Guinea pigs (Cavia porcellus). Journal of Archaeological Science-Reports 17, 678–688. (doi:10.1016/j.jasrep.2017.12.004).

[45] Laffoon, J.E., Valcarcel Rojas, R. & Hofman, C.L. 2013 OXYGEN AND CARBON ISOTOPE ANALYSIS OF HUMAN DENTAL ENAMEL FROM THE CARIBBEAN: IMPLICATIONS FOR INVESTIGATING INDIVIDUAL ORIGINS. Archaeometry 55, 742–765. (doi:10.1111/j.1475-4754.2012.00698.x).

[46] Alscher, S. 2011 Environmental Degradation and Migration on Hispaniola Island. International Migration 49, e164–e188. (doi:10.1111/j.1468-2435.2010.00664.x).

[47] Hooghiemstra, H., Olijhoek, T., Hoogland, M., Prins, M., van Geel, B., Donders, T., Gosling, W. & Hofman, C. 2018 Columbus’ environmental impact in the New World: Land use change in the Yaque River valley, Dominican Republic. Holocene 28, 1818–1835. (doi:10.1177/0959683618788732).

[48] Galloway, J.H. 1996 Botany in the service of empire: The Barbados cane-breeding program and the revival of the Caribbean sugar industry, 1880s-1930s. Annals of the Association of American Geographers 86, 682–706. (doi:10.1111/j.1467-8306.1996.tb01772.x).

[49] Hedges, S.B. & Conn, C.E. 2012 A new skink fauna from Caribbean islands (Squamata, Mabuyidae, Mabuyinae). Zootaxa, 1–+.

[50] Lewis, D.S., van Veen, R. & Wilson, B.S. 2011 Conservation implications of small Indian mongoose (Herpestes auropunctatus) predation in a hotspot within a hotspot: the Hellshire Hills, Jamaica. Biological Invasions 13, 25–33. (doi:10.1007/s10530-010-9781-0).

[51] Schoener, T.W. & Schoener, A. 1983 The time to extinction of a colonizing propagule of lizards increases with island area. Nature 302, 332–334. (doi:10.1038/302332a0).

[52] Gibson, L.M., Mychajliw, A.M., Leon, Y., Rupp, E. & Hadly, E.A. 2019 Using the past to contextualize anthropogenic impacts on the present and future distribution of an endemic Caribbean mammal. Conservation Biology 33, 500–510. (doi:10.1111/cobi.13290).

[53] Kemp, M.E. & Hadly, E.A. 2015 Extinction biases in Quaternary Caribbean lizards. Global Ecology and Biogeography 24, 1281–1289. (doi:10.1111/geb.12366).

[54] Pires, M.M., Guimaraes, P.R., Galetti, M. & Jordano, P. 2018 Pleistocene megafaunal extinctions and the functional loss of long-distance seed-dispersal services. Ecography 41, 153–163. (doi:10.1111/ecog.03163).

[55] Vizentin-Bugoni, J., Tarwater, C.E., Foster, J.T., Drake, D.R., Gleditsch, J.M., Hruska, A.M., Kelley, J.P. & Sperry, J.H. 2019 Structure, spatial dynamics, and stability of novel seed dispersal mutualistic networks in Hawai’i. Science 364, 78–+. (doi:10.1126/science.aau8751).

[56] Barnosky, A.D., Hadly, E.A., Gonzalez, P., Head, J., Polly, P.D., Lawing, A.M., Eronen, J.T., Ackerly, D.D., Alex, K., Biber, E., et al. 2017 Merging paleobiology with conservation biology to guide the future of terrestrial ecosystems. Science 355, eaah4787. (doi:10.1126/science.aah4787).

[57] Jones, H.P., Holmes, N.D., Butchart, S.H.M., Tershy, B.R., Kappes, P.J., Corkery, I., Aguirre-Munoz, A., Armstrong, D.P., Bonnaud, E., Burbidge, A.A., et al. 2016 Invasive mammal eradication on islands results in substantial conservation gains. Proceedings of the National Academy of Sciences of the United States of America 113, 4033–4038. (doi:10.1073/pnas.1521179113).

[58] Holmes, N.D., Spatz, D.R., Oppel, S., Tershy, B., Croll, D.A., Keitt, B., Genovesi, P., Burfield, I.J., Will, D.J., Bond, A.L., et al. 2019 Globally important islands where eradicating invasive mammals will benefit highly threatened vertebrates. Plos One 14, e0212128. (doi:10.1371/journal.pone.0212128).

[59] Fukami, T., Wardle, D.A., Bellingham, P.J., Mulder, C.P.H., Towns, D.R., Yeates, G.W., Bonner, K.I., Durrett, M.S., Grant-Hoffman, M.N. & Williamson, W.M. 2006 Above– and below-ground impacts of introduced predators in seabird-dominated island ecosystems. Ecology Letters 9, 1299–1307. (doi:10.1111/j.1461-0248.2006.00983.x).

[60] Turvey, S.T., Meredith, H.M.R. & Scofield, R.P. 2008 Continued survival of Hispaniolan solenodon Solenodon paradoxus in Haiti. Oryx 42, 611–614. (doi:10.1017/S0030605308001324).

[61] Grant, T.D. & Hudson, R.D. 2015 West Indian iguana Cyclura spp reintroduction and recovery programmes: zoo support and involvement. International Zoo Yearbook 49, 49–55. (doi:10.1111/izy.12078).

[62] Earnhardt, J., Velez-Valentin, J., Valentin, R., Long, S., Lynch, C. & Schowe, K. 2014 The Puerto Rican parrot reintroduction program: Sustainable management of the aviary population. Zoo Biology 33, 89–98. (doi:10.1002/zoo.21109).

[63] Bergstrom, D.M., Lucieer, A., Kiefer, K., Wasley, J., Belbin, L., Pedersen, T.K. & Chown, S.L. 2009 Indirect effects of invasive species removal devastate World Heritage Island. Journal of Applied Ecology 46, 73–81. (doi:10.1111/j.1365-2664.2008.01601.x).

[64] Giovas, C.M. 2019 The Beasts at Large – Perennial Questions and New Paradigms for Caribbean Translocation Research. Part I: Ethnozoogeography of Mammals. Environmental Archaeology 24, 182–198. (doi:10.1080/14614103.2017.1315208).

